# Emergence of social recognition in auditory and integrative circuits during pair bonding

**DOI:** 10.64898/2026.06.01.729418

**Authors:** Erin Wall, Nicholas Vidas-Guscic, Jasmien Orije, Elisabeth Jonckers, Marleen Verhoye, Annemie Van der Linden, Sarah C. Woolley

## Abstract

Social relationships profoundly shape the perception of communication signals. In animals that form long-term social bonds, sensory cues such as vocalizations, scent, or physical appearance, and rewarding mating experiences can become associated with a bonded mate. While this natural process has hallmarks of some forms of associative learning, little is known about where or how these experiences shape neural circuits over the course of pair bonding. We used whole-brain BOLD fMRI in female songbirds to explore experience-dependent changes in neural activity in response to male songs. Pair bonding experiences drove changes in activation in response to the partner’s song within secondary auditory areas as well as two regulatory hubs that receive sensory input and connect to circuits that generate behavioral responses. In contrast, mere exposure to the vocal interactions of a neighboring pair did not result in changes in response to the neighbor’s song within the same areas, suggesting that pair bonding uniquely impacts auditory responses to familiar songs in select sensory and integrative circuits. Moreover, within the areas that showed increased activation for the partner’s song over the course of pair bonding, we observed diminished activation in response to the partner’s song after blocking D1 dopamine receptors, indicating a potential role for dopamine in the expression or maintenance of preference for the partner’s song. Eavesdropping females had changes in neural activation in response to the familiar song in a different suite of brain regions that were not affected by dopamine manipulation. Our data identify a potential circuit in which concurrent activation of sensory, reward, and regulatory hubs integrates multisensory information from social interactions and leads to the emergence of acoustic preferences and formation of long-lasting social bonds.

## Main Text

The motivation to form social bonds is a driving force in social species, including humans. The formation and strength of social attachments depends on dynamic social interactions that shape the perception of sensory cues associated with another individual. While social bonding leads to the learning of sensory cues for individual recognition, social bonds rely not only on sensory learning, but also on preference formation, as a partner’s sensory cues become not just familiar but preferred (*1*–*4*). Pair bonds, defined in part by a preference for a mating partner over other individuals, facilitate breeding coordination and bi-parental care in socially monogamous species (*5*–*11*). For example, coordinated prairie vole parents have offspring that show more parental investment as adults (*9*). In the blue-footed booby, long-term pairs produced more fledglings, suggesting pair bonding enhances reproductive success(*12*).

Zebra finches are opportunistic breeders that form long-lasting pair bonds with a low occurrence of extra-pair copulation (*13*–*16*). Learned songs performed by a male partner are important for social cohesion and bonding, and females rapidly acquire preferences for the song of a male partner with whom they have physically and acoustically interacted (*17*–*19*). Pair-bonded females show differential neural responses in auditory regions and dopaminergic neurons of the ventral tegmental area to the mate’s song compared to an unfamiliar song (*19, 20*). In particular, activity in a secondary auditory region, the caudomedial nidopallium (NCM), hypothesized to be significant in auditory memory, and in the ventral tegmental area (VTA) differs for the mate’s song compared to songs from familiar or unfamiliar males (*20*). While these data highlight the capacity for pair bonding experience to modulate neural responses, the breadth of circuits that have been investigated is limited. The shift from recognition to preference likely requires plasticity in sensory, social, and reward neural circuits highlighting the need for a whole-brain approach for understanding how or where this transformation occurs.

Here, we leveraged the fact that socially monogamous zebra finches rapidly form pair bonds and learned preferences for a partner’s song to investigate the neural representation of a social partner. We took an unbiased, longitudinal approach, using whole-brain BOLD fMRI (blood oxygenation level-dependent functional magnetic resonance imaging) over the course of pair bonding to uncover regions important for the formation and maintenance of partner’s song preferences. This allowed for an unprecedented degree of analysis of brain plasticity in response to social and mating experiences. We also compared the BOLD responses of mated females (females paired with a male partner) listening to the partner’s song to the BOLD responses of unmated females that were housed in same-sex pairs that were eavesdropping on a mated pair to disentangle the contributions of familiarity versus relational identity in driving the neural representations of song. Finally, we used pharmacological manipulations to test the degree to which activation of nodes within the circuit was dependent on local dopamine release. Across a range of species, dopamine (DA) is key for assigning reward value and valence, including in social interactions (*21*–*25*). Dopamine appears to facilitate the formation and expression of partner preferences in zebra finches and populations of dopaminergic neurons in the ventral tegmental area (VTA) increase their activity for preferred songs, including the mate’s song (*20, 26*–*34*). We hypothesized that there would be distinct, experience-dependent changes in brain activation that would differ between mated and eavesdropping birds. For mated females, we further hypothesized that these changes would be modulated by dopamine, as social interaction drives an auditory signal to become not just familiar but also preferred.

## Results

### Pair bonding experience and not song familiarity alone promotes auditory plasticity in the caudal forebrain

Male and female zebra finches form lasting pair bonds and females show learned preferences for the song of their partner. However, the neural circuits underlying this preference, and the extent to which changes in activity within these circuits emerge over the course of preference formation, is unknown. Here, we used non-invasive functional imaging (fMRI) at multiple time points over the course of pair bond formation to investigate the impact of social experience on song perception in the brain. We housed females for two weeks either with a male (mated) or with another female (unmated), separated by an opaque barrier in the same cage as a mated pair. We recorded BOLD activation in mated and unmated females in response to familiar songs of their partner or a neighbor and unfamiliar songs at three time points over two weeks (Figure 1 A,B; see Methods). We first used an unbiased voxel-based analysis that allowed us to examine whether and where changes in activation occur over time in response to song. As has been previously reported, playback of song elicited robust responses throughout the caudal auditory forebrain (*29*). In particular, across all three timepoints and for mated and unmated females, BOLD activation was increased in the primary auditory region Field L and in secondary auditory regions in the caudomedial mesopallium (CMM) and nidopallium (NCM) (p <0.01 FWE) (Figure 1 C,D).

**Fig. 1.**
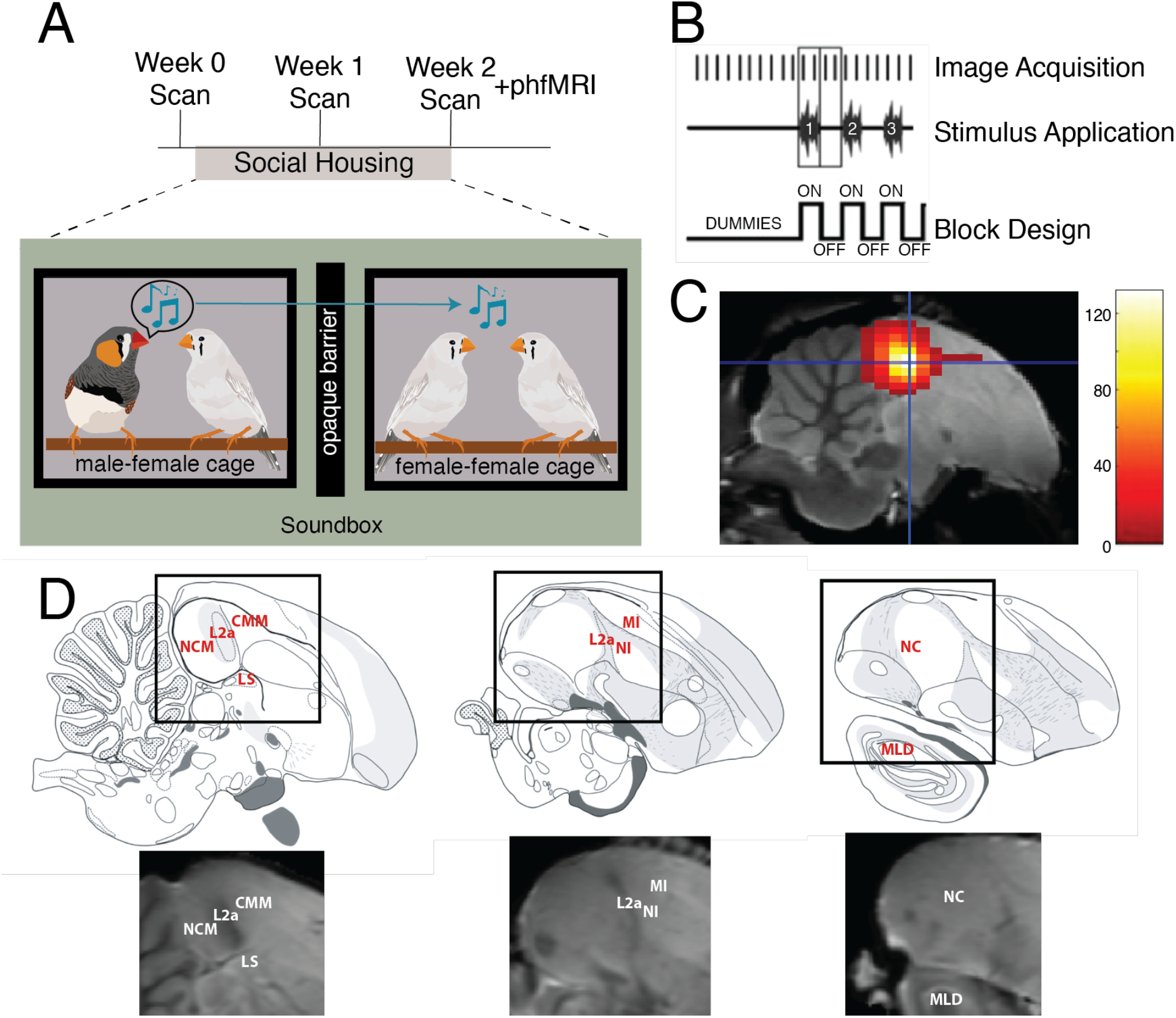
(A) Experimental timeline. fMRI scans were acquired in a 7T MR scanner while females listened to songs. Birds were scanned prior to co-housing (Week 0 Scan) and after one and two weeks of cohabitation. In the cohabitation setup (bottom), a male and female (left, mated) were paired inside a cage (black square) within a sound attenuating chamber (green background). In a neighboring cage (right, unmated) but separated by an opaque barrier, two females were housed together. Within a soundbox, all females could hear the male’s song (blue music notes) but only the female within the same cage could see and physically interact with him. (B) During scans, females heard three categories of song (familiar male’s song, unfamiliar male’s song, and heterospecific male’s song). Images were acquired using ON - OFF block design. During each ON block, two images were acquired during the presentation of a stimulus, followed by an OFF block, where two images were acquired without stimulus presentation. Song stimuli were played in a randomized order throughout a scan. (C) A statistical parametric map of the mean activation in response to all song stimuli for all females in the experiment. Activation in response to song stimuli was greatest in the caudal portion of the pallium, including the auditory forebrain. Crosshairs center on the peak voxel of activation within each cluster. The voxels t-values are color-coded by a heatmap corresponding to the scale on the right. Using this mean activation map, we restricted subsequent analysis to the voxels that responded to auditory stimuli. (D) Sagittal section line drawings adapted from the zebra finch histological atlas (Oregon Health & Science University, Portland, OR 97239; http://www.zebrafinchatlas.org) with a magnified detailed view of the region outlined in black displaying the locations of the labeled areas on the study based template.

While auditory responses were widespread, there were distinct experience-dependent changes in activation that arose over time. We explored how changes in activation vary depending on an individual’s relationship to a song by comparing BOLD activation in response to the familiar song compared to rest at all time points (i.e. before pairing (week 0) and 1 and 2 weeks after pairing) within each subject. Overall, familiar songs (i.e., partner song for mated females and neighbor songs for unmated females) elicited different patterns of activation in mated compared to unmated females. In mated females, four key areas changed significantly in activation over time in response to the familiar song including two regions of the secondary auditory pallium (the CMM and the NCM), as well as two regulatory hubs: the caudocentral nidopallium (NCC) and the lateral septum (LS; Figure 2, Table 1). In the CMM, responses to partner’s song in the left hemisphere increased after females were paired with a male for one week (week 1), and further increased after two weeks of cohabitation (week 2; 2 > 0 and 1, p_uncorr_ = 0.0009; 2 and 1 > 0, p_uncorr_ = 0.0019; Figure 2A,B). In the NCM, a cluster in the ventral part of the NCM showed increased activation in response to the partner’s song following one week of pairing, and remained high at two weeks (2 and 1 > 0, p_uncorr_ = 0.0063; Figure 2C,D). Similarly, a cluster in the LS also showed greater left-hemisphere activation in response to the partner’s song following one and two weeks of cohabitation with a male (2 and 1 > 0, p_uncorr_ = 0.0013; Figure 2 E,F). Lastly, within the NCC, a regulatory hub that has previously been shown to play a role in the evaluating acoustic information important for mate choice (*29*), there were bilateral increases in activation in response to the partner’s song over the two weeks of cohabitation (2 > 0 and 1, Left: p_uncorr_ = 0.0004; Right: p_uncorr_ = 0.0035; 2 and 1 > 0, Right: p_uncorr_ = 0.0014; Figure 2G-J).

**Fig. 2.**
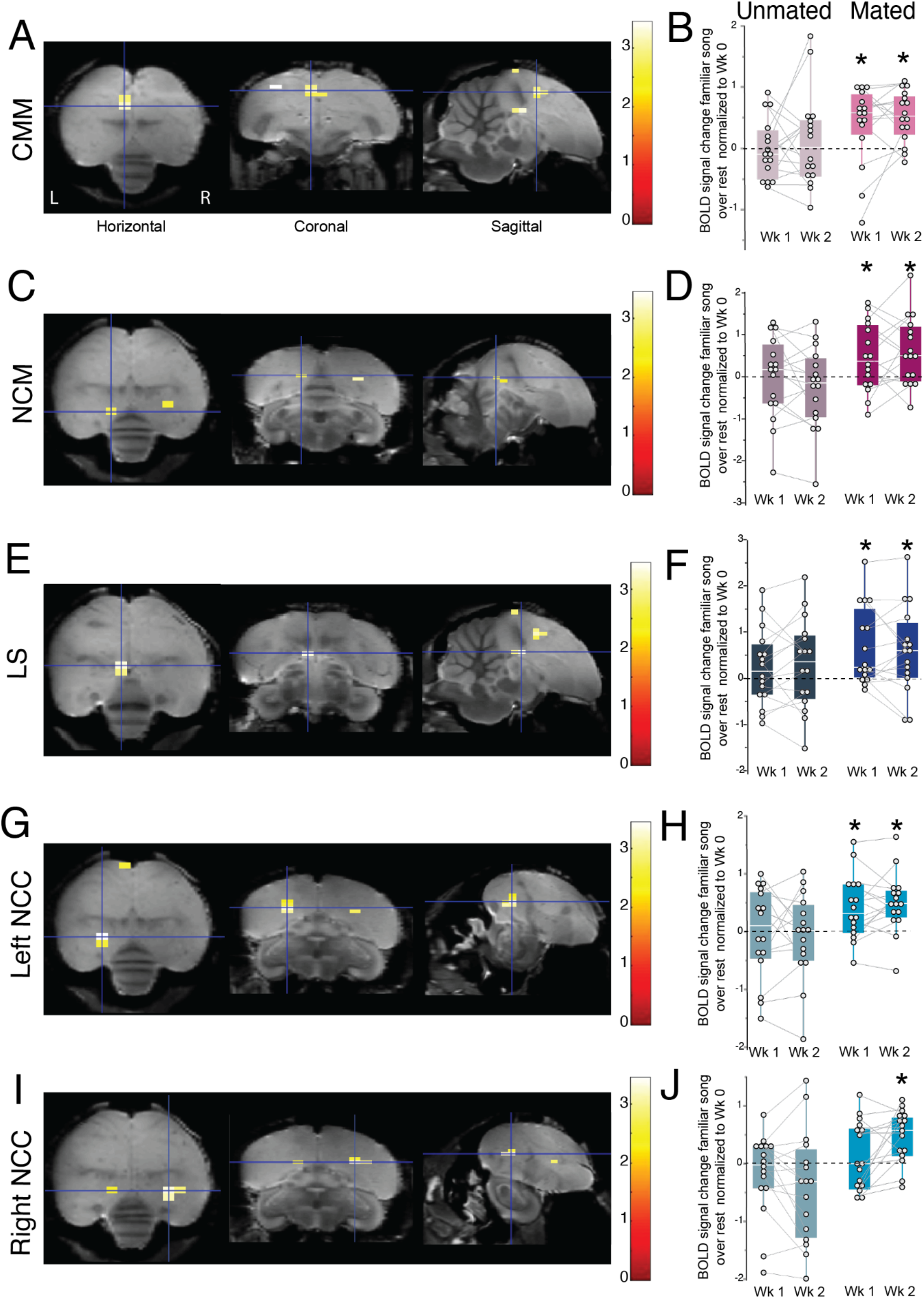
Experience-dependent changes in BOLD responses to the familiar song over rest. (A, C, E, G, I) Statistical parametric maps from voxel-based analysis displaying areas of activation that increase over time for the familiar male’s song over rest (baseline) in mated females (p<0.01). Shown are axial, coronal, and sagittal views (left to right) of the brain using a study-based template from each subject’s 3D anatomical scan. Crosshairs center on the peak voxel of activation within each cluster. (B, D, F, H, J) Box and whisker plots of region of interest based analysis plotted as the BOLD signal change normalized by Week 0 for unmated (left column; light colors) and mated (right column, saturated colors) females on the Week 1 and 2 timepoints. Points are for each individual, boxes span the interquartile range, horizontal white lines indicate the median and whiskers show the minima and maxima. Parametric maps and activation changes over time are shown for the CMM (A,B), NCM (C,D), LS (E,F), Left NCC (G,H) and Right NCC (I, J). Symbols above the box plot indicate the timepoint is significantly different from Week 0, lines and symbols between boxplots indicate a significant differences between Week 1 and Week 2 (* indicates p<0.05; # indicates p<0.11). See Table 1 for cluster sizes, and Table 2 for ROI analysis results.

**Table 1.**
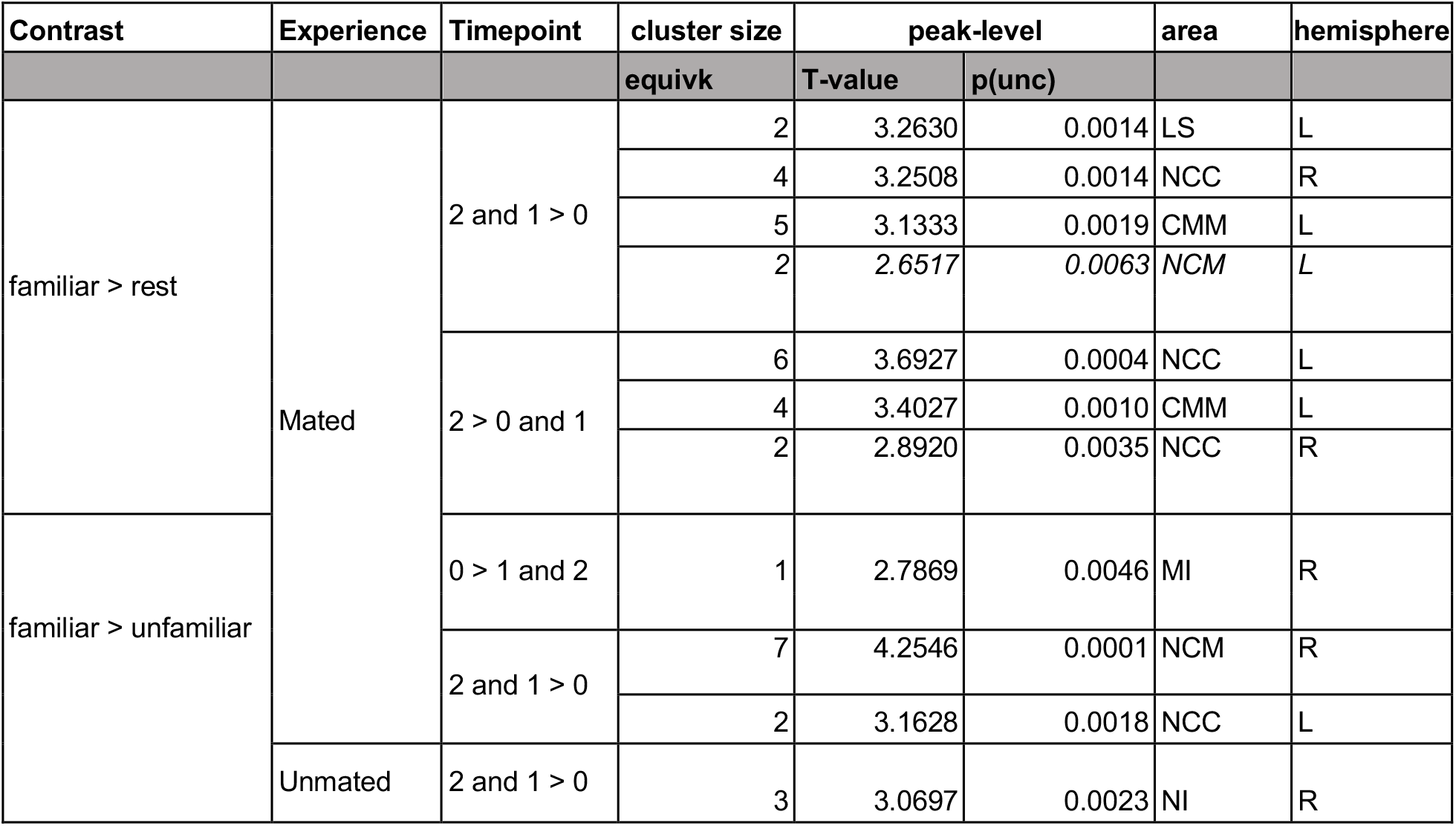
Voxel-based analysis results for activation in response to the familiar song over time. Conditions of interest contrasted for voxel-based analysis (see Figure 2). Stimulus contrast (Contrast) is defined during the first-level analysis for each subject. Timepoint contrast is defined during the second-level analysis to examine changes in activation over time. Cluster size column is the equivk or number of voxels within a cluster at p < 0.01. The t-value represents the T-max for the cluster. The p(unc) states the p-value of the peak voxel in the cluster. We report the voxels that are equal to or less than p < 0.005. Italicized items represent a trend towards significance. Each area is associated with the cluster of activation (see abbreviations list). L - left, R - right hemisphere.

| Contrast | Experience | Timepoint | cluster size | peak-level |  | area | hemisphere |
| --- | --- | --- | --- | --- | --- | --- | --- |
|  |  |  | equivk | T-value | p(unc) |  |  |
| familiar > rest | Mated | 2 and 1 > 0 | 2 | 3.2630 | 0.0014 | LS | L |
|  |  |  | 4 | 3.2508 | 0.0014 | NCC | R |
|  |  |  | 5 | 3.1333 | 0.0019 | CMM | L |
|  |  |  | 2 | 2.6517 | 0.0063 | NCM | L |
|  |  | 2 > 0 and 1 | 6 | 3.6927 | 0.0004 | NCC | L |
|  |  |  | 4 | 3.4027 | 0.0010 | CMM | L |
|  |  |  | 2 | 2.8920 | 0.0035 | NCC | R |
| familiar > unfamiliar |  | 0 > 1 and 2 | 1 | 2.7869 | 0.0046 | MI | R |
|  |  | 2 and 1 > 0 | 7 | 4.2546 | 0.0001 | NCM | R |
|  |  |  | 2 | 3.1628 | 0.0018 | NCC | L |
|  | Unmated | 2 and 1 > 0 | 3 | 3.0697 | 0.0023 | NI | R |

**Table 2.** ROI-based analysis results for activation in response to the familiar song over rest. (see Figure 2). First, we report the area of interest explored using ROI based analysis. The effect reports the main and interaction effects from our statistical analysis. Planned post-hoc comparisons between timepoints are reported by the contrasts with significant results in bold.

| Full Flexible Factorial Analysis |  |  | Planned Post-hoc Comparisons |  |  |
| --- | --- | --- | --- | --- | --- |
| Area | Effect | P-Value | Experience | Contrast | P-Value |
| CMM L | condition x timepoint $F_{2,60} = 2.1632$ | 0.1238 | Mated | 2 vs 0 | **0.0018** |
| | timepoint $F_{2,60} = 3.9569$ | **0.0243** | | 1 vs 0 | **0.0197** |
| | condition $F_{1,30} = 2.2901$ | 0.1407 | | 2 vs 1 | 0.3919 |
|  |  |  | Unmated | 2 vs 0 | 0.4811 |
|  |  |  |  | 1 vs 0 | 0.881 |
|  |  |  |  | 2 vs 1 | 0.3936 |
| NCM L | condition x timepoint $F_{2,60} = 4.0343$ | **0.0227** | Mated | 2 vs 0 | **0.0069** |
| | timepoint $F_{2,60} = 1.1131$ | 0.3352 | | 1 vs 0 | **0.0379** |
| | condition $F_{1,30} = 0.0106$ | 0.9188 | | 2 vs 1 | 0.502 |
|  |  |  | Unmated | 2 vs 0 | 0.2313 |
|  |  |  |  | 1 vs 0 | 0.9006 |
|  |  |  |  | 2 vs 1 | 0.2827 |
| LS L | condition x timepoint $F_{2,60} = 1.1488$ | 0.3239 | Mated | 2 vs 0 | **0.0071** |
| | timepoint $F_{2,60} = 6.0268$ | **0.0041** | | 1 vs 0 | **0.0023** |
| | condition $F_{1,30} = 0.1146$ | 0.7374 | | 2 vs 1 | 0.6984 |
|  |  |  | Unmated | 2 vs 0 | 0.1429 |
|  |  |  |  | 1 vs 0 | 0.2967 |
|  |  |  |  | 2 vs 1 | 0.6674 |
| NCC R | condition x timepoint $F_{2,60} = 3.5027$ | **0.0364** | Mated | 2 vs 0 | **0.0026** |
| | timepoint $F_{2,60} = 2.2109$ | 0.1185 | | 1 vs 0 | **0.0128** |
| | condition $F_{1,30} = 2.0648$ | 0.1611 | | 2 vs 1 | 0.5669 |
|  |  |  | Unmated | 2 vs 0 | 0.6456 |
|  |  |  |  | 1 vs 0 | 0.9126 |
|  |  |  |  | 2 vs 1 | 0.7261 |
| NCC L | condition x timepoint $F_{2,60} = 7.1579$ | **0.0016** | Mated | 2 vs 0 | **0.0059** |
| | timepoint $F_{2,60} = 0.1148$ | 0.8917 | | 1 vs 0 | 0.4035 |
| | condition $F_{1,30} = 1.4561$ | 0.237 | | 2 vs 1 | **0.0484** |
|  |  |  | Unmated | 2 vs 0 | **0.0181** |
|  |  |  |  | 1 vs 0 | 0.2821 |
|  |  |  |  | 2 vs 1 | 0.1837 |

We performed the same unbiased, voxel-based analysis on the scans of the unmated females. In striking contrast to the plasticity in response patterns seen in mated females, we did not find any significant clusters for the familiar song over rest within the caudal forebrain regions that showed increased activation with song playback (for whole brain activation results, see Supplemental Figure S1 and Table S1), indicating that familiarity induced by listening to a neighboring male is not sufficient to drive changes in activation in the caudal forebrain. Rather, social housing with a male seems to dramatically influence neural responses to familiar songs.

The whole-brain, voxel-based analysis allows us to statistically test responses in mated and unmated females separately. In order to directly compare patterns of activation between the social conditions, we next performed a Region-of-Interest (ROI) based analysis focusing on the four regions that were differentially active in mated females in the voxel-based analysis: the CMM, NCM, LS, and NCC (Figure 2). Mated and unmated females show different patterns of activation over time in the NCM and NCC (L and R) (NCM: condition x timepoint F(2,60) = 4.0343, p = 0.0227; Right, condition x timepoint F (2,60) = 3.5027, p = 0.0364; Left, condition x timepoint F (2,60) = 7.1579, p = 0.0016) (Table 2). In the NCC, BOLD activation in mated females increased significantly over the two weeks (planned post-hoc comparison: 2 vs 0 both hemispheres p < 0.01), while in unmated females activation significantly decreased over the same time period (Left, p = 0.0181). In the CMM and LS, BOLD activation increased significantly over the two weeks regardless of condition (CMM: timepoint F(2,60) = 3.9569, p = 0.0243; LS: timepoint F(2,60) = 6.0268, p = 0.0041) (Table 2). Thus, ROI-based analysis further highlighted that pair bonding experiences induce changes in activation in these four forebrain regions, while auditory-only familiarity with a male does not drive these same patterns of activation.

### Mated and unmated females show different patterns of activation changes for the familiar song over unfamiliar song in the caudal forebrain

Our first investigation highlighted regions that showed changes in activity relative to baseline (rest). To assess whether there were areas that respond selectively to the familiar song compared to an unfamiliar song and whether these emerge with social experience, we next explored differential activation in response to familiar vs unfamiliar song using voxel-based analysis (Figure 3).

**Fig. 3.**
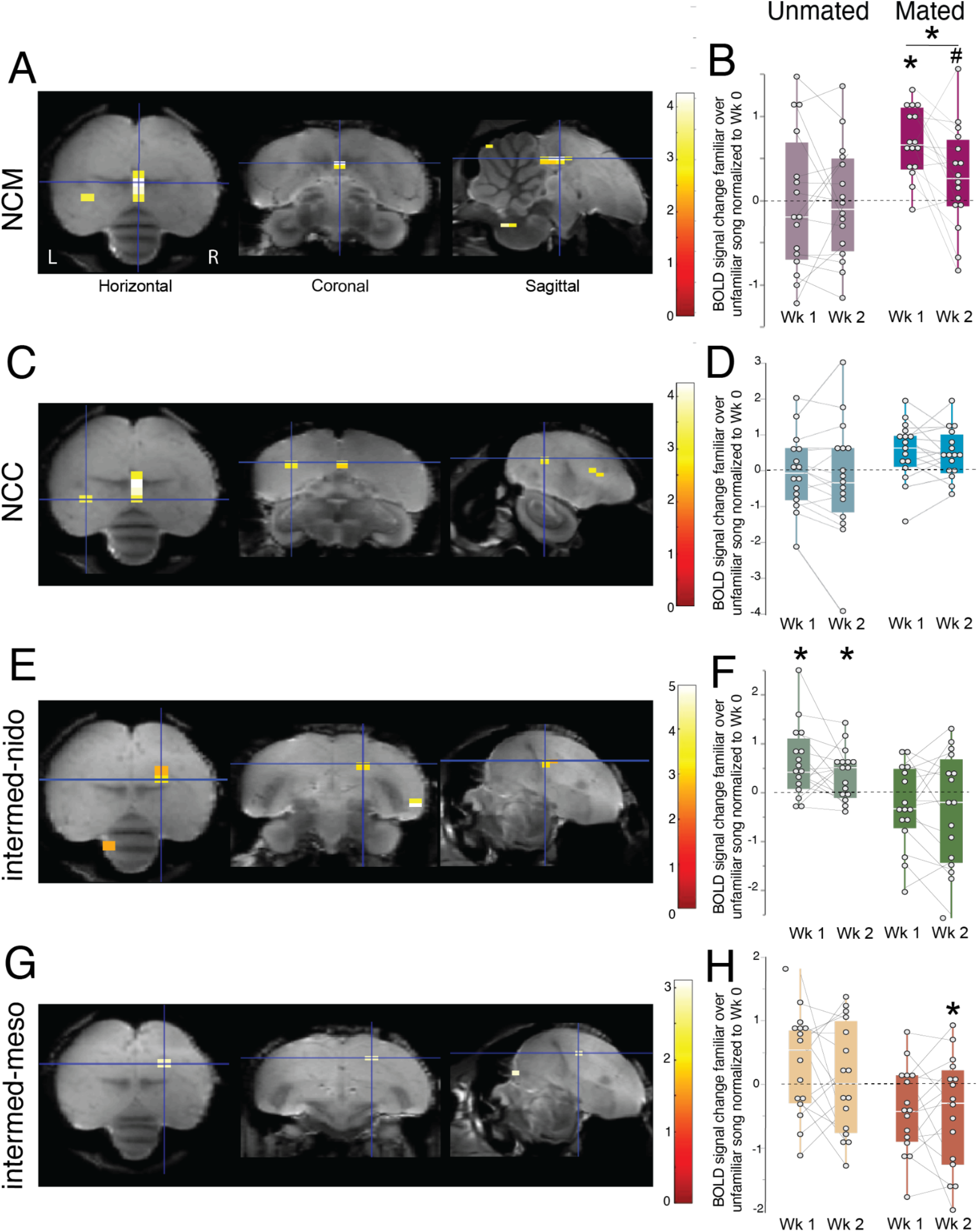
Experience-dependent changes in BOLD responses to familiar over unfamiliar song. (A, C, E, G) Statistical parametric maps from voxel-based analysis displaying areas of activation that increase over time for the familiar male’s song over an unfamiliar male’s song in mated females (p <0.01). Shown are axial, coronal, and sagittal views (left to right) of the brain using a study-based template from each subject’s 3D anatomical scan. Crosshairs center on the peak voxel of activation within each cluster. (B, D, F, H, J) Box and whisker plots of region-of-interest based analysis. Plotted as the BOLD signal change normalized by Week 0 for unmated (left column; light colors) and mated (right column, saturated colors) females on the Week 1 and 2 timepoints. Points are for each individual, boxes span the interquartile range, horizontal white lines indicate the median and whiskers show the minima and maxima. Parametric maps and ROI based activation changes over time are shown for the NCM (A,B), NCC (C,D), NI (E,F), and MI (G,H). * indicates p<0.05; # indicates p<0.15; symbols above the box plot indicate the timepoint is significantly different from Week 0, lines and symbols between boxplots indicate significant differences between Week 1 and Week 2. See Table 1 for cluster sizes, and Table 3 for ROI analysis results.

**Table 3.** ROI- based analysis results in response to the familiar over unfamiliar song over time. (see Figure 3).

| Full Flexible Factorial Analysis |  |  | Planned Post-hoc Comparisons |  |  |
| --- | --- | --- | --- | --- | --- |
| Area | Effect | P-Value | Experience | Contrast | P-Value |
| NCM R | condition x timepoint $F_{2,90} = 4.7042$ | **0.0114** | Mated | 2 vs 0 | 0.1078 |
| | timepoint $F_{2,90} = 3.9110$ | **0.0235** | | 1 vs 0 | **< 0.0001** |
| | condition $F_{1,90} = 0.0237$ | 0.8779 | | 2 vs 1 | **0.0146** |
|  |  |  | Unmated | 2 vs 0 | 0.8309 |
|  |  |  |  | 1 vs 0 | 0.8355 |
|  |  |  |  | 2 vs 1 | 0.9953 |
| NCC L | condition x timepoint $F_{2,89} = 2.2504$ | 0.1113 | Mated | 2 vs 0 | 0.0738 |
| | timepoint $F_{2,89} = 0.7770$ | 0.4629 | | 1 vs 0 | **0.0408** |
| | condition $F_{1,89.4} = 0.3361$ | 0.5636 | | 2 vs 1 | 0.7902 |
|  |  |  | Unmated | 2 vs 0 | 0.3414 |
|  |  |  |  | 1 vs 0 | 0.7546 |
|  |  |  |  | 2 vs 1 | 0.5219 |
| MI R | condition x timepoint $F_{2,90} = 2.7468$ | 0.0695 | Mated | 2 vs 0 | **0.0490** |
| | timepoint $F_{2,90} = 0.8085$ | 0.4487 | | 1 vs 0 | 0.0818 |
| | condition $F_{1,90} = 1.7946$ | 0.1837 | | 2 vs 1 | 0.814 |
|  |  |  | Unmated | 2 vs 0 | 0.7582 |
|  |  |  |  | 1 vs 0 | 0.149 |
|  |  |  |  | 2 vs 1 | 0.2546 |
| NI R | condition x timepoint $F_{2,90} = 4.7684$ | **0.0108** | Mated | 2 vs 0 | 0.1201 |
| | timepoint $F_{2,90} = 0.7994$ | 0.4528 | | 1 vs 0 | 0.2165 |
| | condition $F_{1,90} = 3.7663$ | 0.0554 | | 2 vs 1 | 0.7461 |
|  |  |  | Unmated | 2 vs 0 | 0.0840 |
|  |  |  |  | 1 vs 0 | **0.0051** |
|  |  |  |  | 2 vs 1 | 0.2627 |

In mated females, a ventral area of the NCM (Right) showed higher activation for the familiar partner’s song over an unfamiliar song after pair bonding experience (2 and 1 > 0, p_uncorr_ < .0001) (Figure 3A, Table 1). We also found that the NCC (Left) was significantly more active for familiar song over unfamiliar song after cohabitating with the male partner (2 and 1 > 0, p_uncorr_ = 0.0017) (Figure 3C). In contrast, a cluster near the intermediate mesopallium (MI), a multisensory associative area, showed changes over time in the opposite direction (*35*). In particular, in the MI we found a decrease in activation for the familiar partner’s song over an unfamiliar male’s song over time in mated females (Right, 0 > 1 and 2, p_uncorr_ = 0.00456) (Figure 3G).

Unlike mated females, there were no changes in responses in the NCM, NCC, or MI of unmated females. However, unmated females did show changes to BOLD activation in the intermediate nidopallium (NI), an area involved in imprinting located rostral to the primary auditory area Field L2a (*36*). Activation of this region significantly increased for the familiar partner’s song over unfamiliar male’s song (2 and 1 > 0, p_uncorr_ = 0.0022) in the right hemisphere but was unchanged in mated females (Figure 3E).

We then further investigated activity changes within these four regions (NCM, NCC, MI, NI) using the ROI-based analysis to better directly compare the patterns of activation between mated and unmated females. In the ventral NCM, ROI-based analysis revealed a significant interaction between social condition and timepoint (F(2,90) = 4.7042, p = 0.0114) (Table 3). The difference in activation for mated females was greatest after one week of experience (planned post-hoc comparison p < .0001), followed by a decrease between week 1 and week 2 (p = 0.0146), and a trend towards a difference between week 2 and week 0 (p=0.1078) (Figure 3B, Table 3). Unmated females did not show any significant differences between timepoints (p > 0.8 for all timepoint comparisons). In the left NCC, although we found increased activity for the familiar song over unfamiliar song in only mated females in the voxel-based analysis, we did not find significant effects of condition, timepoint, or the interaction in the ROI-based analysis (Figure 3D, Table 3). In the MI, ROI-based analysis found a trend towards an interaction effect (F(2,90) = 2.7468, p = 0.0695) with a significant decrease in activation between week 0 and week 2 in mated females (p = 0.0490) (Figure 3H, Table 3). Finally, for the NI cluster, there was a significant interaction of condition and timepoint (F(2,90) = 4.7684; p = 0.0108), as only unmated females showed a significant change in activation over time (week 1 vs 0 p = 0.0051) (Figure 3F, Table 3). Thus, changes in BOLD activation over the two weeks occurs in both mated and unmated females, but in different regions.

### A dopamine receptor antagonist diminishes mate’s song responses

Several of the pallial regions captured by our voxel-based analyses, including the NCM, NCC, and LS, are heavily innervated by dopaminergic terminals and express dopamine receptors, particularly D1 receptors (*27, 30*). For example, the NCM receives dopaminergic input from the ventral tegmental area (VTA) and substantia nigra pars compacta (SNc), and a dopamine D1 agonist acting locally in the NCM can alter females’ song preferences (*26*). We hypothesized that the changes in activity we see in response to a male’s song as females form pair bonds with a partner may be driven or maintained by dopamine. To explore the role of dopamine in the response to familiar song following two weeks of pair-bonding, we scanned females for an additional session in Week 2 (pharmacological fMRI: phfMRI) after peripheral injection of either a D1 receptor antagonist or saline. The phfMRI scan included the same set of stimuli as the Week 2 scan, allowing for comparison of pre- and post-pharmacological manipulation (see Methods).

We investigated where changes in activation in response to the familiar song occurred pre vs post manipulation for both social (mated and unmated) and drug (antagonist and saline) conditions using a voxel-based analysis. Among mated females, D1 receptor antagonism led to decreased activation for the familiar song vs rest in three areas: NCM (Left, p_uncorr_ = 0.0085), LS (Left, p_uncorr_ = 0.0039), and NCC (Right, p_uncorr_ = 0.0054) (Table 4; Figure 4). Strikingly, both the LS and NCC clusters are the same as (or within one voxel of) those that were significantly more active for the familiar song over rest before drug manipulation (LS, Left: p_uncorr_ = 0.0026; NCC, Right: p_uncorr_ = 0.0012) (Figure 4G). Such changes in responses were not observed in mated females treated with saline. Similarly, activation of these clusters did not change in unmated females treated with either the D1 antagonist or saline (no suprathreshold clusters p_uncorr_ < 0.01). These results suggest that bonding experience led to greater activation of dopaminergic inputs in response to the partner’s song.

**Table 4.**
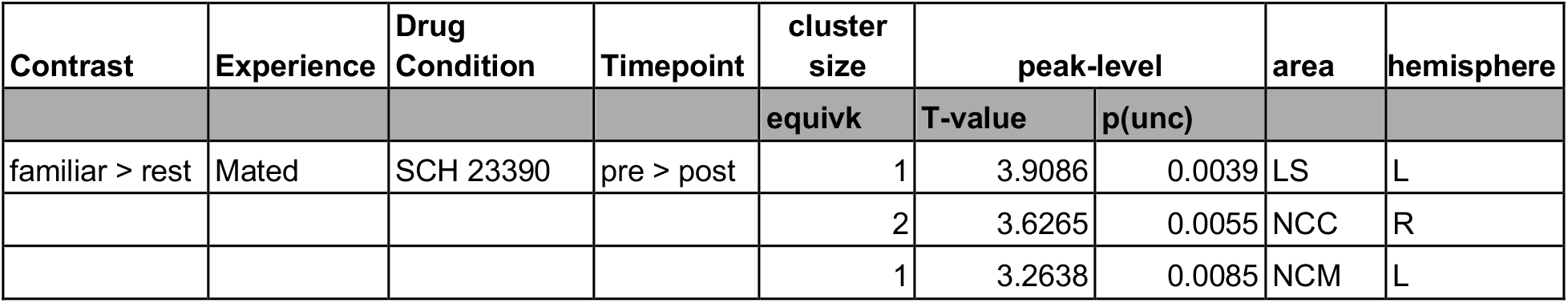
Results table for the voxel-based analysis of phfMRI (see Figure 4). Mated females show significant decreases in activation in response to the familiar song.

**Fig. 4.**
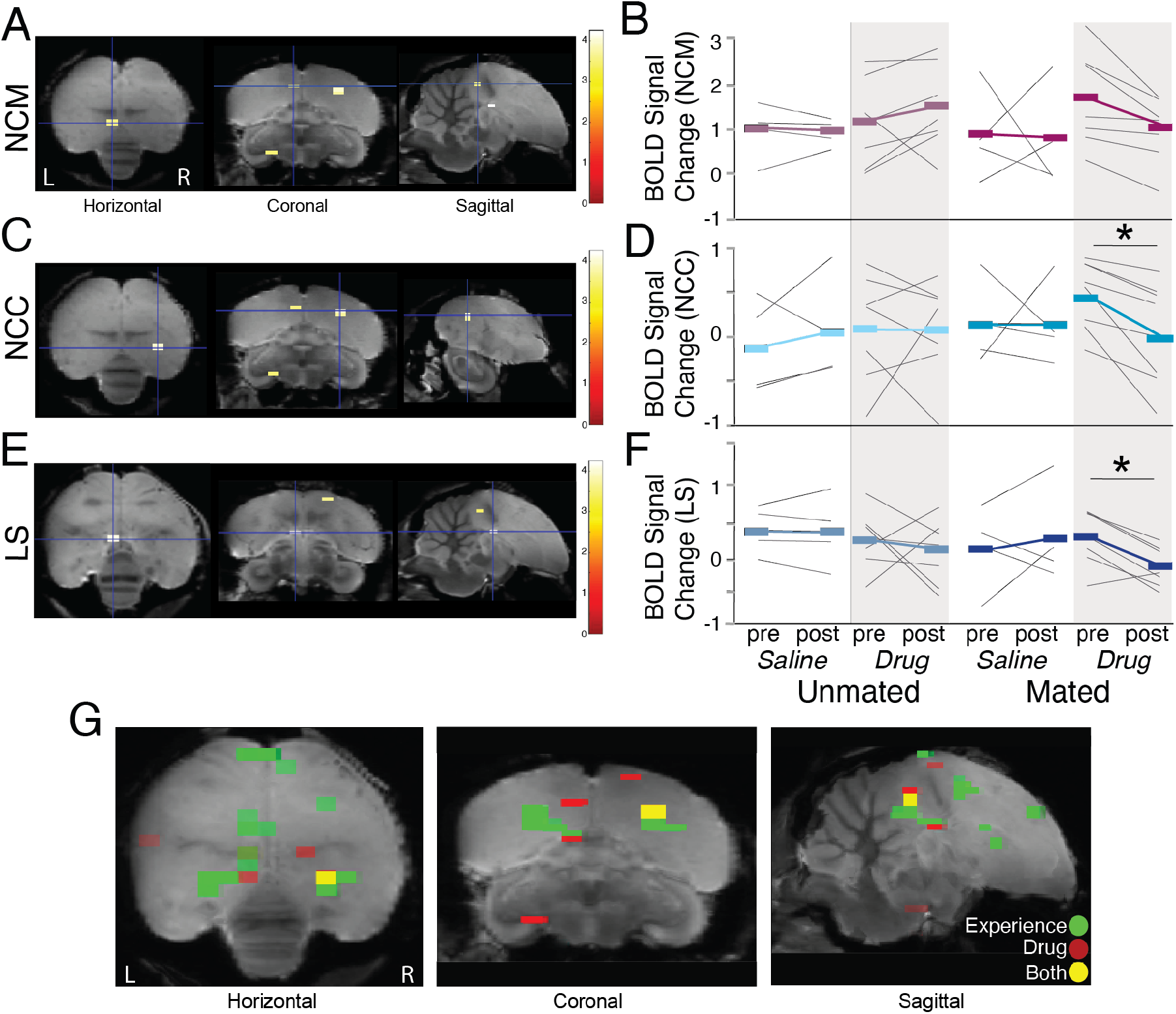
A D1 antagonist diminishes BOLD responses in regions that show changes with pair bonding. (A,C,D) Voxel-based analysis for the changes in activation before vs. after D1 antagonist administration in mated females in the NCM (A), NCC (C), and LS (E) (p<0.01). (B,D,F) ROI-based analysis displaying the pattern of activation pre vs post pharmacological manipulation (saline or D1 antagonist), measured by the change in BOLD signal, in unmated (left) and mated females (right) treated with either saline (white shading) or drug (D1 antagonist; gray shading) in the NCM (B; magenta), NCC (D; cyan), and LS (F; blue). Colored bars and lines indicate the mean, thin lines are responses of individual birds. (G) Map of the proximity of clusters of activation. Clusters of activation that increase over time in response to the familiar song (over rest) for mated females are shown in green (“experience”) and the clusters that show a decrease in activation following dopamine manipulation with a D1 antagonist are shown in red (“drug”). Voxels that both increase over time and decrease with drug manipulation are shown in yellow. Images are overlaid and extend from midline to ∼2.5mm lateral. * indicates p<0.05. See Table 4 for cluster sizes, and Table 5 for ROI analysis results.

**Table 5.** Results table for the ROI-based analysis of phfMRI (see Figure 4). The effect of condition refers to the experience condition, mated or unmated. The effect of timepoint refers to the week 2 timepoint (pre-drug or saline) and the phfMRI scan (post-drug or saline). The effect of drug refers to the drug condition, saline or SCH22390.

| Full Flexible Factorial Analysis |  |  | Planned Post-hoc Comparisons |  |  |  |
| --- | --- | --- | --- | --- | --- | --- |
| Area | Effect | P-Value | Experience | Drug Condition | Contrast | P-Value |
| NCM L | condition x timepoint x drug $F_{1,18} = 1.3766$ | 0.256 | Mated | SCH23390 | pre vs post | 0.0647 |
| | timepoint x drug $F_{1,18} = 0.0620$ | 0.8062 | | saline | pre vs post | 0.8231 |
| | conditon x drug $F_{1,18} = 0.0420$ | 0.84 | Unmated | SCH23390 | pre vs post | 0.3396 |
| | condition x timepoint $F_{1,18} = 1.7991$ | 0.1965 | | saline | pre vs post | 0.9388 |
| | drug $F_{1,18} = 1.5992$ | 0.2221 | | | | |
| | timepoint $F_{1,18} = 0.3513$ | 0.5608 | | | | |
| | condition $F_{1,18} = 0.0209$ | 0.8866 | | | | |
| LS L | condition x timepoint x drug $F_{1,18} = 0.1474$ | 0.7056 | Mated | SCH23390 | pre vs post | **0.0394** |
| | timepoint x drug $F_{1,18} = 4.6623$, | **0.0446** | | saline | post vs pre | 0.5724 |
| | condition x drug $F_{1,18} = 0.8233$ | 0.3762 | Unmated | SCH23390 | pre vs post | 0.3272 |
| | condition x timepoint $F_{1,18} = 0.4247$ | 0.5228 | | saline | pre vs post | 0.4272 |
| | drug $F_{1,18} = 0.1984$ | 0.6613 | | | | |
| | timepoint $F_{1,18} = 0.3529$ | 0.5599 | | | | |
| | condition $F_{1,18} = 5.4716$ | **0.0311** | | | | |
| NCC R | condition x timepoint x drug $F_{1,18} = 0.3583$ | 0.5569 | Mated | SCH23390 | pre vs post | **0.0379** |
| | timepoint x drug $F_{1,18} = 1.7187$ | 0.2063 | | saline | pre vs post | 0.9988 |
| | condition x drug $F_{1,18} = 0.0254$ | 0.8751 | Unmated | SCH23390 | pre vs post | 0.9623 |
| | condition x timepoint $F_{1,18} = 1.6175$ | 0.2196 | | saline | pre vs post | 0.559 |
| | drug $F_{1,18} = 0.1818$ | 0.6749 | | | | |
| | timepoint $F_{1,18} = 0.4109$ | 0.5296 | | | | |
| | condition $F_{1,18} = 0.4281$ | 0.5212 | | | | |

We performed an ROI-based analysis to directly compare between drug and social conditions within each of these three clusters. In the LS cluster, we found a significant interaction between time point and drug condition (F(1,18) = 4.6623, p = 0.0446) (Figure 4, Table 5). Mated females that received the D1 receptor antagonist showed significantly higher responses to the familiar song before pharmacological manipulation (planned post-hoc comparison p = 0.0394) while females that received saline and unmated females treated with either drug or saline did not show this change in activation (p > 0.3). In the NCC, activation also decreased, and trended towards a decrease in NCM, following the D1 receptor antagonist in mated females (planned post-hoc comparison NCC: p = 0.0379; NCM p = 0.0647); however, when comparing this subset of females to the other conditions, there were no significant effects of condition (NCC: F(1,18) = 0.4281, p = 0.5212; NCM: F(1,18) = 0.0209, p = 0.8866), timepoint (NCC: F(1,18) = 0.4109, p = 0.5296; NCM: F(1,18) = 0.3513, p = 0.5608), drug condition (NCC: F(1,18) = 0.1818, p = 0.6749; NCM: F(1,18) = 1.5992, p = 0.2221), or the interaction (NCC: F(1,18) = 0.3583, p = 0.5569; NCM: F(1,18) = 1.3766, p = 0.256) (Figure 4, Table 5).

## Discussion

During the formation of stable, long-term pair bonds, salient social interactions lead to a memory of and a preference for the sensory signals associated with a partner. We found that female finches given the opportunity to socially interact and bond with a male exhibited stronger activation in two auditory areas, the NCM and CMM, and two regulatory hubs, the NCC and LS, in response to their partner’s song after two weeks of cohabitation. In contrast, females that had the same acoustic exposure as mated females but could not see or physically interact with the male showed changes in activation in a completely distinct complement of brain regions. Moreover, after two weeks of pair bonding, blocking dopamine D1 receptors led to a decrease in activation in the NCM, NCC, and LS in mated but not unmated females, indicating that hearing the partner’s song leads to local release of dopamine into these regions in bonded females. Taken together, these changes in brain activation for a newly familiar, highly salient song highlight how the nature of a social relationship impacts the perception of auditory signals. Further, these data lend insight into the neural pathways and modulators involved in learned recognition and preference that are fundamental to communication, social networks, and reproduction.

The auditory cortex has been implicated as a key node for auditory learning and plasticity, and storage of auditory memories (*37*–*40*). In line with previous work, our results demonstrate that courtship songs elicit activity in the CMM and that this response is enhanced by experience (*29, 41, 42*). The NCM, in contrast, is hypothesized to encode song memories that have alternately been thought to be represented as increased selectivity or as increased habituation to songs from specific individuals such as the tutor or the mate(*19, 20, 43*–*45*). Intriguingly, our results revealed both increased activation for a partner’s song over unfamiliar song and, in the dorsal NCM, habituation with repeated exposure(*43, 44, 46*–*48*). The increases and decreases appeared in distinct locations, hinting that there may be region-specific variation in the encoding or processing of song within NCM.

Our voxel and ROI based approaches uncovered two hub regions, the NCC and LS, which, based on their connections and patterns of activity, have the potential to link social, sensory, and reward information necessary for bonding and preference formation. The diverse connections of the NCC to sensory, learning and memory, and mesolimbic reward circuitry, and to hypothalamic and motor areas that coordinate behavioral performance suggest that the NCC could serve as an interface for the processing of and response to salient sensory stimuli (*29*). Across vertebrates, the LS is a part of a social decision-making network (SDMN) that regulates social behavior. In mammals, the LS, is linked to social affiliation, recognition, reproduction, and parental care, regulating affect and behavioral outputs (*49*–*51*). In songbirds, the limited data on septal function has focused on male zebra finches where LS activity is increased during nest building and vasotocin release into the LS modulates gregariousness and anxiety (*52, 53*). Our data, which are the first to indicate a potential role of the LS in pair bonding in females finches, bolster support for a similar role of the LS in social behavior of monogamous birds as has been described in monogamous mammals.

The LS, NCM, and NCC are each heavily innervated by catecholamines and express dopamine receptors (*27, 29, 30, 54*). Dopaminergic projections from regions including the ventral tegmental area (VTA) and substantia nigra (SNc) are key to evaluating the valence of sensory signals, ascribing salience to sensory stimuli, and shaping social decision-making (*23, 34, 55*– *58*). Moreover, dopaminergic inputs to the auditory cortex can also drive sensory plasticity and manipulation of dopamine can modify or enhance auditory perception and preference (*26, 32, 55, 56, 59, 60*). To test whether activation in response to the partner’s song was modulated by dopamine, we coupled pharmacological manipulation using a dopamine antagonist together with whole-brain imaging. We found diminished activity to the partner’s song in the NCM, NCC, and LS following injection of a D1 antagonist, but only in mated females, and not in any saline treated females. These data indicate that hearing the partner’s song drives dopamine release into these regions following two weeks of pair bonding. Interestingly, both the NCC and LS clusters neighbor or overlap with areas that increased in activation over time in mated females, suggesting that dopamine plays a role in shaping the response to partner’s song in these areas (Figure 4G) and highlighting a role for dopamine in driving the plastic changes we observe over pair bonding in auditory and sensory integration hubs.

Communication facilitates the transfer of information, mediates social decisions, and drives social behavior, making it critical to survival in many species. Perception is shaped by experience throughout life, and therefore requires finely-tuned and plastic neural circuitry for processing the content, context, and salience of vocal signals. The perception of song is critical for females to make mate choice decisions, recognize familiar conspecifics, and maintain a life- long pair bond with a mate. Based on the results presented here, we propose a potential model for auditory preference from sensation to action, integrating areas of auditory processing, reward, and social decision-making, whereby females evaluate vocal signals, form preferences, and respond (Figure 5). In mated females, we speculate that secondary auditory areas (NCM, CMM) encode information about the acoustic properties of song and singer identity. Concurrently, activation of reward areas such as VTA or nucleus accumbens (NAcc) attribute valence to the acoustic signals. Together, the NCC and the LS integrate multisensory information, including coupling song with physical interactions from social bonding experiences to promote preference learning. The LS and NCC, potentially via motor regions such as the medial arcopallium, relay signals to hypothalamus and dorsomedial nucleus of the midbrain (DM) to drive vocal behaviors, such as callbacks and reproductive behaviors such as copulation solicitation displays. Taken together, our data build a framework for understanding how experience influences perception and provide the foundations of circuitry important for preference learning and partner recognition.

**Fig. 5.**
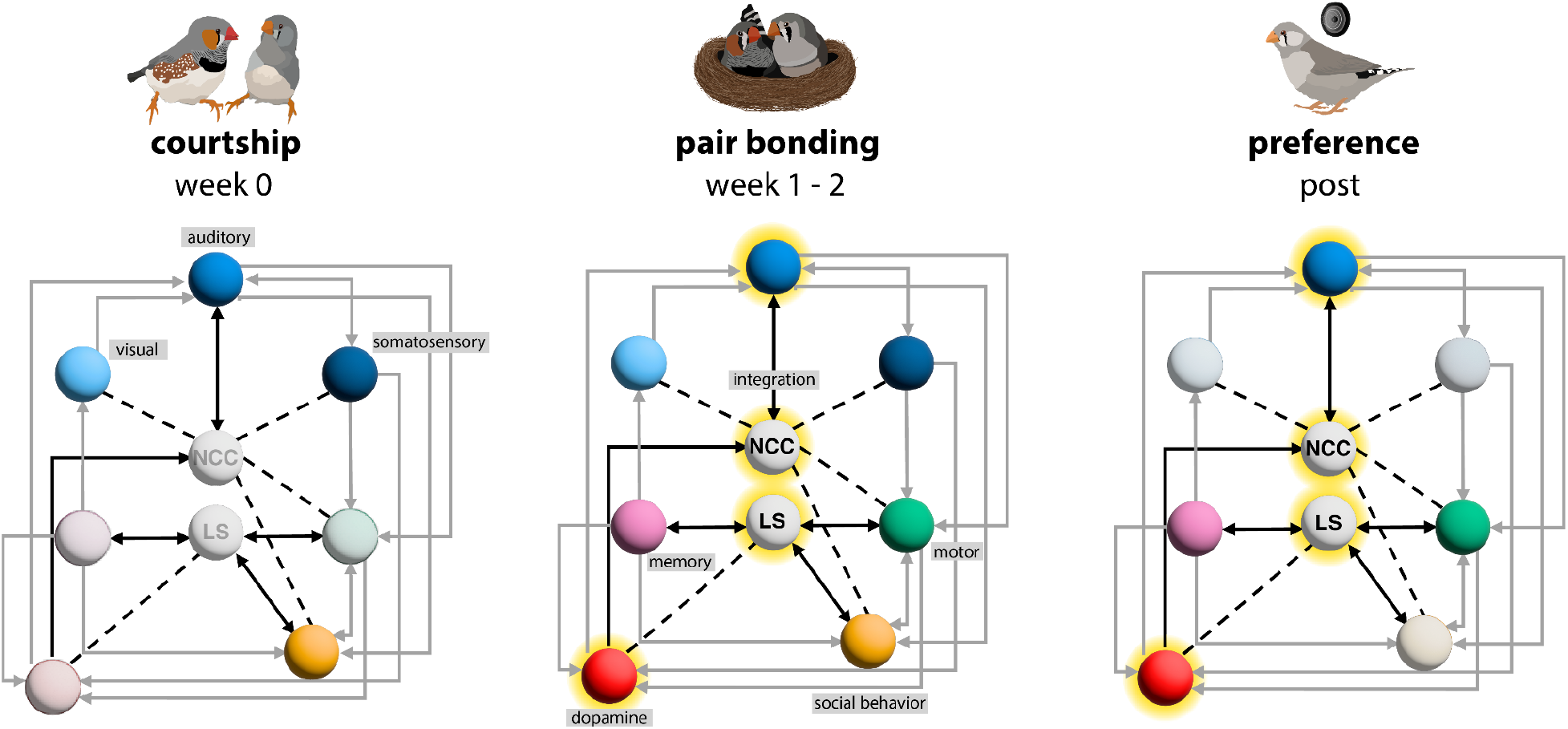
A proposed model of neural activity changes during social bonding. Each panel depicts a network of regions (by their associated function) hypothesized to be involved at each timepoint. Solid grey lines represent connections between regions in zebra finches. Solid black lines highlight connections with the proposed integration hubs, NCC and LS. Dotted lines represent connections found in other model species, or connections with the NCL in zebra finches that are not specific to the NCC. The saturation of the nodes reflects the hypothesized degree of involvement for the respective experience. Highlighted nodes (glow) indicate areas that showed a change in activity within this study.

## Acknowledgements

We sincerely thank Johan Van Auderkerke for guidance and support throughout the acquisition and analysis of the data, and Christien Bowman for assistance with data collection and methodology. We’d also like to thank Julie Hamaide for guidance and collaboration during the preparation of the study. In addition, we thank Constance Scharff and her lab at the Freie Universität Berlin for providing many of the birds used in this study and for meaningful discussion about the experiments. We are grateful for the insights and technical support from Tamara Vasilkovska, Monica van den Berg, Maarten Naeyaert, Gaurav Majumdar, Verdi Van Reusel, Lisbeth Van Ruijssevelt, and all members of the Bio-Imaging Lab small animal imaging core facility where the experiment was conducted. Thank you to Jon T. Sakata and Clementine Bodin for helpful discussions and comments on the data analysis and results.

## Supplemental Materials

Materials and Methods

Abbreviations

Tables 1-5

Supplementary Text

Table S1

Figure S1

## Materials and Methods

### Subjects

Normally-reared, adult female zebra finches without prior mating experience (n = 32, > 90 days old) were used for the experiments at the Bio-Imaging Lab at the University of Antwerp (Antwerp, Belgium). Zebra finches were either bred at the Bio-Imaging Lab facilities (University of Antwerp, Antwerp, Belgium, n = 8 females, n = 9 males), acquired from the animal colony of a colleague (Constance Scharff, Freie Universität Berlin, n = 20 females), or purchased from a registered breeder (n = 4 females, n = 7 males). All zebra finches were housed in large same-sex group aviaries (males and females kept in separate rooms) on a 12 hr light/dark cycle and given access to food, water, and sand/grit *ad libitum* and given weekly supplements (i.e. egg, greens). All procedures and animal handling were performed in accordance with the European guidelines for the care and use of laboratory animals (2010/63/EEC) and approved by the Committee on Animal Care and Use at the University of Antwerp, Belgium (ECD 2019–17). At the end of this non-invasive neuroimaging study, the birds were returned to the aviaries of the University of Antwerp.

### Cohabitation

Prior to the Week 0 timepoint, all females were housed in same-sex group cages in a separate room from all males. We then manipulated the social experiences of females by housing them in one of two conditions for two weeks (Figure 1A) (*18*). After each subject’s Week 0 scanning session, once they were fully recovered from the anesthesia, females were allocated to their cohabitation conditions (Figure 1A). Each female was housed either with one of the conspecific males whose song she heard during the Week 0 scan (opposite-sex pair: mated) or with a female (same-sex pair: unmated). One mated pair and one unmated pair were placed inside a sound-attenuating chamber in a cage subdivided in two, separated by an opaque barrier (dimensions: (105 × 70 × 57) cm^3^). Females sharing the same sound-attenuating chamber are referred to as a “cohort” that share the same auditory experience with an individual male.

After the initial introduction, the sole male in the sound-attenuating chamber is considered familiar to all three females in a cohort. All females in a cohort could hear the male’s songs and vocalizations but only the female housed in the same cage (mated) could see and physically interact with the male. Unmated females only hear the male’s song but do not see or physically interact with him. Each cohort was filmed for at least two hours upon the first introduction, after one week of cohabitating, and after two weeks of cohabitating (see also Wall and Woolley, 2024)(*18*).

#### Auditory stimuli

### Song recordings

Courtship and non-courtship song stimuli were collected as previously described (*19, 41*). Briefly, males’ songs were recorded individually in a sound-attenuating chamber with an omnidirectional microphone (Beringer) using sound activated recording software (44.1 kHz, Sound Analysis Pro) (*61*). To collect courtship songs, females not involved in the experiment housed in a separate cage were presented to a male for brief periods (∼30 seconds).

### Stimulus paradigm

We selected 10 to 15 courtship and non-courtship songs for each male that were a representative sample of the male’s song and free of noise or female calls. All songs were bandpass filtered (300-10kHz) and normalized by maximum amplitude using custom code in Matlab (Mathworks, Natick, MA, USA). Songs were then concatenated in Praat (http://www.praat.org/, University of Amsterdam) to 16 seconds of song with minimal silence between motifs followed by 16 seconds of silence to adhere to the stimulus presentation requirements in an ON/OFF block paradigm for fMRI (Figure 1B). Next, these stimuli were batch processed to match loudness and equalized using a custom setting that counters fMRI frequency distortion in the scanner (Audition, Adobe) (*62*). We adapted the established auditory stimuli preparation protocol of Van Ruijssevelt et al (2013) and assessed the frequency equalization using a fiber-optic microphone (Optimic 1160, Optoacoustics) placed in the bore during pilot testing (*62*).

Three sets of song stimuli were presented during each fMRI scan: two conspecific males’ songs and one heterospecific (Bengalese finches, recorded at McGill University) male’s songs. For each scanning session, we acquired two separate fMRI scans for each subject: a courtship song scan, containing the courtship songs of the three stimulus types, and a non-courtship song scan, containing the non-courtship songs of the three stimulus types. In this study, we focused our second-level analyses on the conspecific courtship song scans. During the first scanning timepoint, Week 0, all stimuli were unfamiliar to all subjects. After the scanning session, females from a cohort were introduced to one of the conspecific males they heard during scanning. This male became the familiar male for that cohort. His songs were played for that cohort during each scan, representing the familiar stimulus type. At each timepoint, the song of a novel conspecific male was used as the unfamiliar stimulus. For each cohort, the two sets of conspecific males’ songs had similar song durations and number of motifs.

#### Functional MRI

We used a 7-Tesla horizontal MR system (Pharmascan 70/16 US, Bruker Biospin, Germany) to image the head of 32 adult female zebra finches in four imaging sessions: Week 0 (before pairing), Week 1 (1 week after pairing), Week 2 (2 weeks after pairing), and phfMRI (2 weeks after pairing).

### Animal Preparation

Prior to scanning, each subject was transferred to a cage in the scanning room. Food and water were removed 40 minutes to 1 hour prior to scanning. As two birds were scanned in a day, three birds were transferred to the scanning room together to minimize social isolation other than during scanning (mated female, unmated female, and companion female). All subjects were anesthetized with isoflurane (2.5% induction; 1.2% maintenance; IsoFlo, Abbott, Illinois, USA) in oxygen and nitrogen (flow rate 100 and 200cm^3^/min, respectively). Birds were placed in a custom-made scanner bed with an adaptable beak mask to provide isoflurane anesthesia and to offer a reproducible positioning of the head for each subject. The setup for auditory fMRI consisted of a volume RF coil for transmission, and a custom-made single loop radio-frequency surface receive (coil) which is tightly placed around the bird’s head. Auditory stimuli were played through the two non-magnetic dynamic speakers (Visation, Germany), placed on either side of the zebra finch head, at 70dB. The acoustic noise created by the switching of the magnetic field gradients of the scanner was decreased by increasing gradients’ ramp time to 1000 µs. From the start of the procedure, body temperature and breathing rate were monitored with a pressure sensitive pad and cloacal thermistor probe (MR-compatible Small Animal Monitoring and Gating System, SA Instruments). To ensure the birds’ body temperature remained stable (at 40.0 ± 0.5 °C), a feedback-controlled warm air circuitry was connected to the scanner bed (MR-compatible Small Animal Heating System, SA Instruments, Inc.). After scanning, birds recovered from anesthesia in a cage with companions under a heating lamp. When subjects fully recovered from anesthesia, they were returned to their cohabitation cages.

### Data acquisition

To quickly determine the correct position of the subject in the scanner, three orthogonal gradient echo images were acquired (referred to as the localizer scan). Once the subject was correctly centered in the scanner, we acquired three multislice Turbo rapid acquisition relaxation-enhanced (RARE) T2-weighted scans in the axial, coronal, and sagittal direction to guide the position of the slice package of the subsequent fMRI scans in a reproducible manner. For each fMRI scan, we acquired 298 image volumes using a T2-weighted (RARE) sequence during stimulus presentation with the following settings: TEeff /TR: 60/2000 ms; RARE factor 8; 15 sagittal slices with the slice thickness 0.8mm and interslice gap distance 0.05mm (encapsulating the entire brain); FOV (16 × 16)mm^2^ ; matrix [64 × 32]; spatial resolution: (0.25 × 0.5) mm^2^; temporal resolution 8 s; total scan duration: 40 min. Each subject also underwent an anatomical 3D T2-weighted RARE scan in the same orientation as the fMRI scans at one of the three timepoints with the following settings: TE 11 ms (TE eff 44 ms), TR 3000 ms, RARE factor 8, FOV (16 × 14 × 14) mm^3^, matrix [256 × 92 × 64], spatial resolution (0.063 × 0.152 × 0.22) mm^3^, scan duration 37 min.

At each timepoint, two fMRI time series scans were performed for each subject while they were in the scanner, constituting a scanning “session.” Females were presented with the courtship songs in one scan, and the non-courtship songs in the other scan. The order of scans was randomized across groups. In this study, we investigated changes in activation over time using the courtship song scans. Each scan presented the three different stimulus types in a randomized order across an ON/OFF block paradigm alternating 16 seconds of stimulation (ON) and 16 seconds of rest (OFF) (Figure 1B). The OFF blocks serve as baseline or activation at rest during the scanner noise without stimulus presentation. In this way, the response to a stimulus can be measured by the statistical difference in signal intensity between the ON and OFF block images. Within each block, two whole-brain images are acquired (8 second scan time). In total, this results in 288 images acquired from 72 ON blocks and 72 OFF blocks, with the first 10 images acquired without stimulus presentation (298 scans in total). Each stimulus type therefore has 48 image volumes per scan.

### fMRI with pharmacological modulation

At the end of the cohabitation period and immediately following the Week 2 fMRI scanning session, we performed a pharmacological fMRI (phfMRI) scan with 22 out of the 32 subjects(*63*). Just after collecting the Week 2 fMRI scan, females were briefly removed from the scanner bed and given a subcutaneous injection (50 µl) into the inguinal fold of either a dopamine receptor (D1) antagonist (SCH23390, 0.3mg/ml; 1mg/kg or 0.001mg/g body weight) (n = 14) or 0.9% saline (n = 8). Care was taken to minimize the time out of the scanner bed to perform the injection (<3 minutes). Following the injection, we performed the localizer and slice package alignment as described above (∼ 30 minutes). Then, a second fMRI scan using stimuli identical to the Week 2 courtship fMRI scan was acquired. The Week 2 courtship fMRI scans before and following the pharmacological manipulation are referred to as the pre and post fMRI scan, respectively, and constitute the phfMRI scanning session.

### Data processing

Image processing and voxel-based statistical analysis was performed using SPM 12 (The Wellcome Centre for Human Neuroimaging, London, UK; https://www.fil.ion.ucl.ac.uk/spm) following protocols adapted from Van Ruijssevelt et al. (2013, 2018) (*29, 62*). To correct for head movement and possible drift, we first realigned all fMRI scans to the first scan using rigid transformation. We inspected the movement parameters for each fMRI scan time series to ensure head motion did not exceed 2 voxels displacement (direction dependent, 0.5 / 1 / 1.6mm translation) or excessive rotation in any direction. Then, we co-registered each subjects’ fMRI scans to their own anatomical 3D RARE. We spatially normalized each subject’s 3D RARE to the study-based template created from all subjects’ 3D RARE images in ANTs. We applied the combined transformation of the coregistration and normalization transformation to bring each subject’s fMRI in the template space. We visually inspected the outcome of the normalization and coregistration steps by comparing the registration of each scan to the study-based template. The voxels measured in the present study have not been upsampled, neither at the level of reconstruction using zero-filling, as this procedure is susceptible to Gibbs-ringing artifacts, nor at the level of image-upsampling (*64*). The voxel size used in these analyses is (0.25 × 0.5 × 0.8) mm^3^; thus, one voxel sampled in our analysis encompasses sixteen “upsampled” voxels (0.125 × 0.125 × 0.400) mm^3^ used in Van Ruijssevelt et al. (2018). We proceeded to smooth the fMRI images in-plane with a (0.5mm × 1mm) Gaussian kernel full width at half maximum (FWHM).

At the first level within-subjects analysis, we modeled the BOLD responses of each subject as a box-car convolved with a canonical hemodynamic response function and estimated the model parameters using a general linear model with Restricted Maximum Likelihood. The movement parameters estimated during the realignment step were included as regressors in the model. We used a high pass filter (352s) to remove signal drift and a whole brain explicit mask so that only the voxels within the brain are analyzed.

#### Statistical Analyses

### Subject-level analysis to test for stimulus-specific activation

To examine differences in brain activation corresponding to the different stimuli, we created - for each subject and time point - t-contrast images for each stimulus over rest and comparisons between stimuli, including familiar vs rest and familiar vs unfamiliar song. We assessed whether each subject’s scans displayed a bilateral positive BOLD response in the auditory forebrain for the stimuli presented. The single subject contrasts between each stimulus > rest were then used in further second-level group analyses.

### Group-level analysis for activation changes over time

First, we explored the mean group activation in response to song at the whole-brain level by including all subjects, all stimuli (familiar>rest vs unfamiliar>rest, and all timepoints in a full flexible factorial analysis. Activation in response to song vs rest was found primarily in the caudal forebrain (cluster size = 492 voxels) at a statistical significance threshold of p_FWE_ <0.01 (Figure 1C). Using the statistical parametric map from this test, we created a mean activation mask to further investigate activation changes in only areas that respond to auditory stimuli. In small animal fMRI, multiple comparison correction such as family-wise error correction (FWE) can be too conservative to detect effects (*29, 63*). For subsequent voxel-based analysis, data were considered significant at p_uncorr_ < 0.005 and reported by the highest voxel T-value within a cluster (Tmax) and the voxel p-value. While the present study examined the areas that showed the greatest change in activation with pair bonding experience within the clusters that are significantly responsive to auditory stimuli compared to rest (FWE <0.01), further insights can be gained from this data set through exploration of all the areas that responded to the auditory stimuli at each timepoint.

First, to examine temporal changes in activation in response to the familiar song with experience, we performed within-subjects repeated measures ANOVAs for mated and unmated females separately, using the single subject t-contrast images for familiar > rest from each timepoint as input. Secondly, in a separate analysis of the single subject t-contrast images for familiar > unfamiliar at each timepoint into a separate analysis, we explored temporal changes in activation where the response to the familiar male’s song was greater than to an unfamiliar male’s song. For each of these within-subject ANOVAs on the t-contrast familiar > unfamiliar, we assessed both the positive and negative change in activation difference between timepoints using the following t-tests: between Week 0 and Week 1, between Week 0 and Week 2, and between Week 1 and Week 2.

To further explore changes in activation over time within-subjects and between social conditions, we performed a Region of Interest (ROI) based analysis. ROIs were selected based on the significant clusters in the group-level voxel based analysis of both familiar vs rest and familiar vs unfamiliar responses (using a significance level of p_uncorr_ < 0.01 to examine more of the region). We used MarsBar (https://marsbar-toolbox.github.io/) to extract for each ROI and at each timepoint the ROI-averaged BOLD contrast from the single subject t-contrast images for familiar > rest and familiar > unfamiliar for all subjects at each timepoint. We performed full flexible factorial analyses with condition and timepoint as factors, BOLD contrast as the dependent variable, and bird ID as a random factor. To compare two timepoints within a condition, we used LSMeans student’s t-test with *α* < 0.05 unless otherwise noted.

### Group-level analysis for change in activation differences with dopamine manipulation

For the second-level voxel-based analysis of activation before and after systemic injection of a dopamine antagonist, we ran a separate flexible factorial analysis with condition (mated and unmated), timepoint (pre vs post), and drug condition (saline and SCH 23390). We used the single subject t-contrast images from the courtship song scan at the week 2 timepoint (pre) and the phfMRI scan (post) for mated and unmated females treated with either drug or saline. To explore where changes in activation differences may occur across the whole brain, we used a whole-brain mask and a statistical threshold of p_uncorr_ < 0.01. We performed paired t-tests to detect differences pre vs post drug manipulation for each condition and drug condition, for both familiar> rest and familiar > unfamiliar. To compare between drug conditions and social conditions, we extracted data from the significant clusters as ROIs using MarsBar as described above. For the purposes of this study, we focused on significant clusters in the caudal forebrain. Next, we performed full flexible factorial analyses for familiar > rest and familiar > unfamiliar, with condition, drug condition, and timepoint as factors, BOLD contrast as the dependent variable, and bird ID as a random factor. To investigate the difference in BOLD response at the level of the ROI within each condition, we also performed LSMeans student’s t-tests with *α* < 0.05. All statistical comparisons were conducted in SPM (12) and JMP (SAS, 16pro).

## Abbreviations

Mated: opposite-sex paired
Unmated: same-sex paired
BOLD fMRI: blood oxygen level dependent functional magnetic resonance imaging
PhfMRI: pharmacological fMRI
RARE: rapid acquisition relaxation-enhanced
ROI: region of interest
AI: intermediate arcopallium
AM: medial arcopallium
AMV (or TnA): ventro-medial arcopallium or nucleus taenia of the amygdala
AV: ventral acropallium
CLM: caudolateral mesopallium
CMM: caudomedial mesopallium
DM: dorsomedial nucleus of the midbrain
Loc: locus coeruleus
LS: lateral septum
MD: dorsomedial mesopallium
MI: intermediate mesopallium
NAcc: nucleus accumbens
NCC: caudocentral nidopallium
NCL: caudolateral nidopallium
NCM: caudomedial nidopallium
NI: intermediate nidopallium
NIL: intermediate lateral nidopallium
NIM: intermediate medial nidopallium
PoA: posterior pallial amygdala
SNc: substantia nigra
VTA: ventral tegmental area

