## Supplemental Text, Table S1, Figure S1 for "Emergence of social recognition in auditory and integrative circuits during pair bonding"

### Supplemental Materials

While our voxel-based analysis using a mean activation mask to examine areas that respond to song stimuli did not find any significant clusters of activation that change over time in unmated females, whole-brain analysis without any explicit activation mask revealed two regions that decreased in activation over time: a dorsal region of the NCM (Figure S1A) and a dorsal region of the CM (lateral to CMM) (Table S1). Interestingly, a dorsal region of the NCM also decreased over time for mated females (T max = 2.615368, p = 0.006884) (Figure S1B). ROI-based analysis also found a significant effect of timepoint driven by the mated females (F2,60 = 3.8116, p = 0.0276) (mated 0 vs 1 p =0.0107, 0 vs 2 p = 0.0220; unmated 0 vs 2 p =0.3415).

###### *Table S1. Unmated females show activation changes in two areas of caudal pallium.*

*In unmated females, whole-brain voxel-based analysis (without FWE < 0.01 mean activation mask) revealed two areas within the caudal forebrain that decrease in activation over time in response to the familiar song: a dorsal area of the caudal nidopallium and a dorsal area of the caudal mesopallium.*

| **Contrast** | **Experience** | **Timepoint** | **cluster size** | **peak-level** | | **area** | **hemisphere** |
| --- | --- | --- | --- | --- | --- | --- | --- |
|  |  |  | **equivk** | **T-value** | **p(unc)** |  |  |
| familiar > rest | Unmated | 0 > 1 and 2 | 5 | 3.8061 | 0.0003 | dNC | L |
|  |  |  | 1 | 2.5713 | 0.0076 | dCM | R |

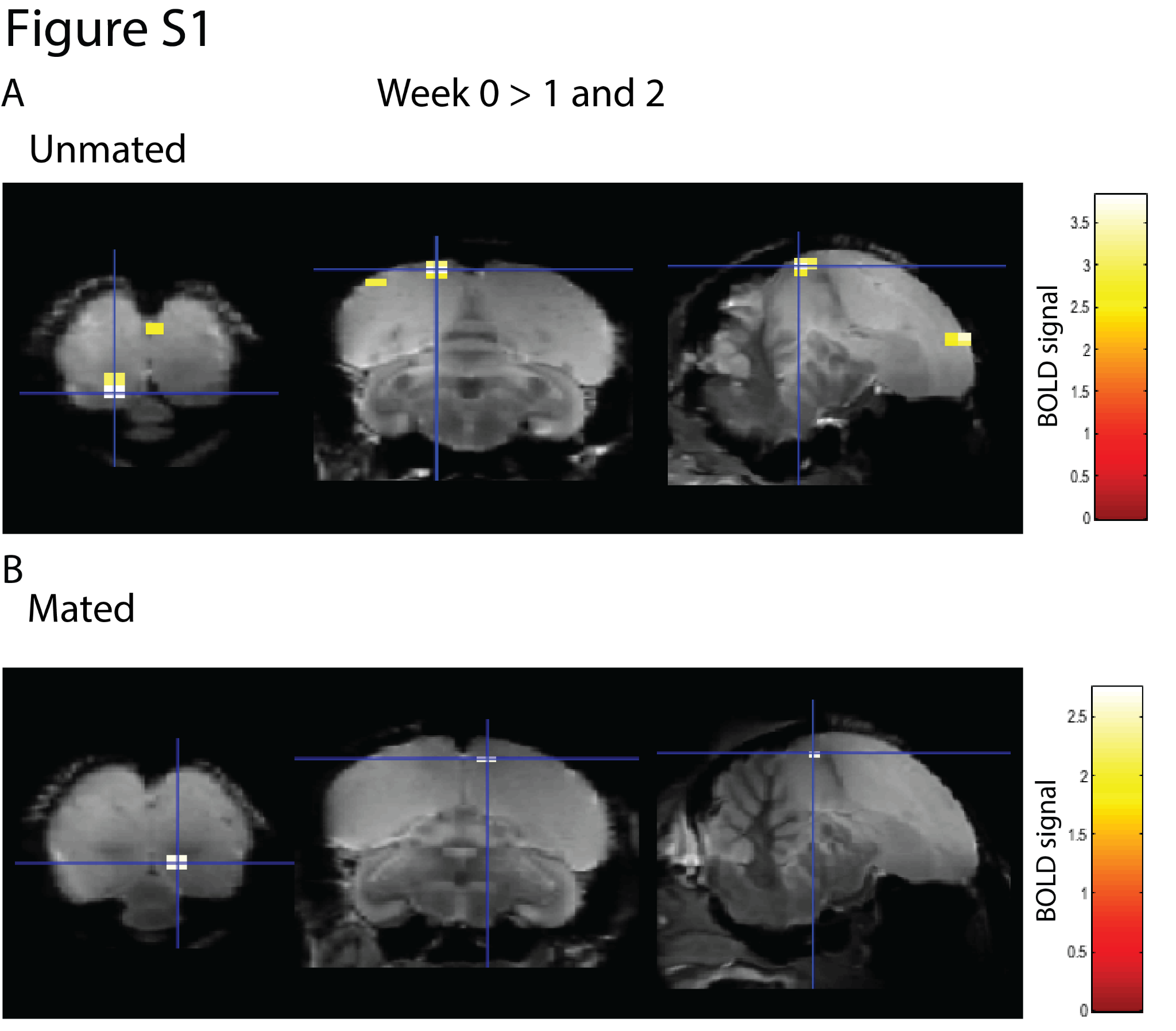

###### *Figure S1. A dorsal region of the caudal nidopallium shows decreased activation over time for the familiar song in both unmated (A) (T = 3.81, p = 0.0003) and mated females (B) (T = 2.62, p = 0.0068).*
